# Bioorthogonal Tethering of FGF18 to Annealed Microgel Scaffolds Enhances Cartilage Repair

**DOI:** 10.64898/2026.08.04.742809

**Authors:** Erika E. Wheeler, Hye-Jeong Jang, Kelly C. Weldon, Kun Chen, Yuting Wang, Katherine H. Griffin, Thomas H. Ambrosi, J. Kent Leach

## Abstract

Articular cartilage damage often progresses to osteoarthritis (OA), a degenerative joint disease characterized by chronic pain, limited mobility, and reduced quality of life. Tissue engineering approaches using poly (ethylene glycol) (PEG)-based hydrogels offer tunable mechanical properties and bioactive functionalization, yet the influence of surface charge on cartilage regeneration remains underexplored. Moreover, recombinant fibroblast growth factor 18 (FGF18) has successfully improved cartilage tissue thickness in clinical trials, but required high dosages and recurring injections may limit compliance. Here, we developed a granular microgel-based platform with bioorthogonally tethered FGF18 to evaluate the interplay of microgel surface charge and growth factor presentation on chondrogenesis. Azide groups were incorporated onto the microgel surfaces to enable site specific FGF18 conjugation across microgel scaffolds with distinct surface charges. When seeded with mesenchymal stromal cells, microgel scaffolds functionalized with FGF18 outperformed their unmodified counterparts, evidenced by higher GAG content, collagen content, and compressive modulus. In a murine microfracture model, anionic and zwitterionic microgel scaffolds tethered with FGF18 increased cartilage regeneration compared to nonionic microgels. We detected increased collagen II content within defects treated with tethered FGF18 microgel scaffolds. This work demonstrates the role of surface charge and growth factor presentation in directing cell behavior and tissue repair, advancing the design of biomaterials for cartilage regeneration.

**HIGHLIGHTS:**

- Covalent tethering of FGF18 to microgel surface enables localized bioactivity
- FGF18 tethered microgel scaffolds enhance extracellular matrix deposition and chondrogenic differentiation of murine mesenchymal stromal cells in vitro
- Localized FGF18 presentation improves cartilage tissue formation and mechanical properties in vivo, with anionic and zwitterionic microgel scaffolds outperforming nonionic scaffolds
- FGF18 presentation is a more potent stimulus than microgel surface charge for cartilage regeneration

## INTRODUCTION

Articular cartilage has a limited capacity for self-repair due to its avascularity and low cell density[1]. Consequently, cartilage degeneration can progress to osteoarthritis (OA), a degenerative joint disease that affects more than 595 million people globally with direct medical costs up to 72 billion dollars in the USA[2, 3]. OA is characterized by chronic pain, impaired mobility, and progressive tissue degeneration[2]. No treatments are currently available that reverse or slow disease progression; therefore, existing therapies are limited to symptomatic relief[4].

Naturally derived polymers such as agarose[5], alginate[6, 7], collagen[8, 9], chitosan[10], and hyaluronic acid[11, 12] have been widely investigated for cartilage regeneration due to their structural similarity to native extracellular matrix (ECM) and inherent bioactivity[13]. However, their use is restricted by poor mechanical stability, batch-to-batch variability, and limited tunability. Synthetic polymers, particularly poly(ethylene glycol) (PEG), offer well-defined chemistry, tunable mechanical properties, and resistance to nonspecific protein adsorption[14]. PEG-based hydrogels can be readily functionalized with bioactive ligands[15, 16], degradable crosslinkers[17, 18], or growth factors[19, 20] to modulate cellular behavior and establish instructive regenerative microenvironments conducive to cartilage tissue regeneration. However, bulk hydrogels are limited in tissue engineering, as the nanoporous mesh must be degraded for cells to proliferate and migrate, and mechanical properties are often related to porosity.

Annealed microgel scaffolds represent a versatile biomaterial platform due to their injectability, modularity, and inherent macroporosity[21]. These scaffolds are composed of individual hydrogel microparticles, commonly formed using microfluidic or emulsion-based methods, that can be annealed into interconnected porous networks using enzymes or light that facilitate rapid cell infiltration, nutrient diffusion, and ECM remodeling[22, 23]. Zwitterionic granular hydrogels enable chondrocyte proliferation, migration, and ECM production[24]. We recently demonstrated that zwitterionic microgel scaffolds support ECM accumulation in inflammatory conditions, and that substrate charge influences protein adsorption, mesenchymal stomal cell (MSC) adhesion, and subsequent ECM deposition[25]. Moreover, pH-responsive zwitterionic nanoparticles presenting kartogenin to damaged cartilage suppressed osteophyte formation and promoted regeneration of collagen II and aggrecan[26]. Despite these advances, the role of microgel surface charge in regulation protein adsorption, immune interaction, and cartilage regeneration remains poorly understood.

Fibroblast growth factor 18 (FGF18) is a clinically relevant anabolic growth factor for articular cartilage regeneration due to its ability to promote chondrocyte proliferation, ECM deposition, and cartilage maintenance through the activation of the FGFR3 signaling pathway[27]. The FORWARD Phase II clinical trial demonstrated that intra-articular injection of human recombinant FGF18 (sprifermin) increased cartilage thickness in patients with OA[28]. Patients receiving the 100-µg dose every six or twelve months exhibited increased cartilage thickness compared to those receiving the 30-µg dose or placebo at two years, although the clinical significance of this increase was not determined. Despite these promising anabolic effects, soluble delivery of FGF18 is constrained by rapid diffusion from the joint space and the requirement for repeated intra-articular injections to maintain therapeutic effect. The sustained delivery of localized growth factors using engineered biomaterials may improve therapeutic efficacy while reducing injection burden.

We incorporated azide groups onto PEG to covalently tether FGF18 on microgels with distinct surface charges. We first characterized the effects of azide incorporation and protein tethering on scaffold properties including zeta potential, protein adsorption, and compressive modulus. We then evaluated the chondrogenic potential of tethered FGF18 within annealed microgel scaffolds of varying surface charge. Finally, we tested the therapeutic potential of this platform in a murine microfracture model and evaluated cartilage regeneration. Collectively, this work establishes a microgel platform for investigating how surface charge and localized growth factor delivery promote cartilage regeneration.

## METHODS

### Fabrication of microgels and growth factor tethering

Microgels were fabricated using an emulsion method in a microfluidic device and crosslinked as previously described[25]. Microgels were formed using poly(ethylene glycol) methyl ether methacrylate (Nonionic, Sigma-Adrich, St. Louis, MO, USA), 3-sulfopropyl methacrylate (Anionic, Sigma-Aldrich), and [2-(methacryloyloxy) ethyl] dimethyl-(3-sulfopropyl) ammonium hydroxide] (Zwitterionic, Sigma-Aldrich). Microgel precursor solutions contained a 10% w/v monomer concentration, 5% w/v of poly(ethylene glycol) dimethacrylate (PEGDMA, MW 750, Sigma-Aldrich), 1 mM Acryl-RGD (GenScript, Piscataway, NJ, USA), and 1 mM of Acryl-PEG-Azide (PegWorks, Chapel Hill, NC, USA) in 25 mM HEPES buffer at pH 7.4 with 0.45% of photoinitiator 2,2’-Azobis[2-methyl-N-(2-hydroxyethyl) propionamide] (VA-086, FUJIFILM, Tokyo, Japan). Microgels were fabricated in a microfluidic device and collected in a glass vial containing 10 mL of mineral oil and exposed to UV light (Omnicure S2000, 20 mW/cm^2^, 320-500 nm, St. Louis, MO, USA) for durations between 2 to 3 min to achieve similar microgel stiffness.

Human recombinant FGF18 (Peprotech, Cranbury, NJ, USA) and bovine serum albumin (Sigma Aldrich, St. Louis, MO, USA) were diluted in sterile 1x PBS at pH 7.25 with 0.1% BSA to create a 1 mg/mL solution. DBCO-FGF18 and DBCO-BSA were synthesized by adding DBCO-ester (Vector Laboratories, Newark, CA, USA) in DMSO at a 1:8 molar ratio to solution of FGF18 and BSA. The reaction took place overnight at 4°C and was quenched using sterile 10x TRIS for 1h at 4°C. The DBCO-FGF18 was stored at -20°C until use. To tether DBCO-FGF18 to microgels, DBCO-FGF18 was diluted in 1% BSA to achieve a concentration of 2 μg/mL once added to the microgels. The DBCO-FGF18 solution was added to the microgel slurry at a 1:1 volume ratio. The azide-microgel/DBCO-FGF18 solution was incubated at 37°C for 1 h prior to annealing. The DBCO-FGF18 solution was aspirated from the microgels, washed with sterile HEPES, and then replaced with annealing solution. To confirm tethering, 30 µM of DBCO-488 staining buffer was added and incubated at 37°C for 35 min. Samples were washed twice with 1% BSA washing buffer.

### Quantification of FGF18 release from microgels

To assess FGF18 release from tethered microgels, we cultured acellular microgel scaffolds in serum-free DMEM for 7 days. Experimental groups included microgels tethered with FGF18 and anionic microgels with soluble FGF18. On day 7, we collected 400 µL of conditioned medium and quantified human FGF18 using a sandwich enzyme-linked immunosorbent assay (ELISA) kit according to the manufacturer’s protocol (Antibodies.com, St. Louis, MO, USA). Due to sample absorbance values approaching the lower limit of the assay range, we calculated FGF18 concentrations by interpolation from a sigmoidal standard curve in GraphPad Prism.

### Surface charge characterization

The zeta potential of precursor solutions was measured using a Zetasizer (Nano ZS90, Malvern, UK). All monomers were resuspended at 10% w/v in DI water and sonicated briefly to prevent aggregation. Samples were loaded into cuvettes, and zeta potential was measured three times. The conductive properties of hydrogels were measured using a previously published custom two-point setup[29].

Acellular microgel scaffolds were stored in 1X PBS, and protein adsorption studies were performed as previously described[25]. After aspirating the storage solution, 1 mL of 40 μg/mL lysozyme (EMD, Darmstadt, Germany) was added and allowed to incubate at 37°C for 24 h. After incubation in lysozyme, the solution in each well was collected and analyzed for protein content using the bicinchoninic acid (BCA) Protein Assay Kit (Pierce, Rockford, IL, USA).

### Measurement of microgel diameter and mechanical testing

Microgels were pipetted onto a petri dish with a glass bottom and imaged at 10x using an EVOS microscope. Microgel diameter was manually measured using ImageJ with at least 3 fields of view. The compressive moduli of individual microgels was determined using a MicroTester (CellScale, Waterloo, ON) by loading onto an anvil in a water bath filled with PBS at pH 7.25[30]. Microgels were compressed over 30 s to 20% of their original diameter by a 2×2 mm stainless-steel platen attached to a 0.2032 diameter tungsten rod. The displacement and force of each microgel was recorded and used to calculate the modulus. The modulus was calculated from the linear region of the compressive modulus versus the nominal strain graph using a custom-made Python code.

### Annealing of microgels into a continuous scaffold

Microgels were annealed as previously described[25]. Briefly, microgels were pelleted at 2000*xg* for 5 min and resuspended in equal volume of annealing solution consisting of PEG-DMA crosslinker in culture medium containing 0.05% (Lithium phenyl-2,4,6-trimethyl-benzoylphosphinate, Sigma). After incubating for 1 min, the microgel solution was jammed *via* centrifuge at 16000*xg* for 3 min. The supernatant was removed and the microgel slurry was optionally mixed with cells.

### MSC cell culture and chondrogenesis

Murine bone marrow-derived mesenchymal stromal cells (mMSCs, C3H10T1/2, ATCC, Manassas, VA, USA) at passage 12-14 were cultured in growth media containing DMEM, 10% fetal bovine serum (FBS), and 1% penicillin/streptomycin (P/S) under standard culture conditions. For chondrogenesis experiments, the microgel slurry was mixed with cells seeded at 5×10^6^ cells per mL and pipetted into 5 mm x 1.5 mm cylindrical molds using a positive displacement pipette (Gilson, Middleton, WI, USA). The microgel slurries were then exposed to UV light (20 mW/cm^2^, 320-500 nm, Omnicure S2000, St. Louis, MO, USA) for 2 min to form cylindrical annealed scaffolds *via* free radical polymerization. Scaffolds were kept in growth media for 24 h after fabrication before switching to chondrogenic medium for 21 d at 21% O_2_, and media changes were performed every other day. Chondrogenic medium was composed of DMEM with 1% P/S, 1.5 mg/mL bovine serum albumin, 100 μg/mL sodium pyruvate, 40 μg/mL L-proline, 50 μg/mL L-ascorbic acid 2-phosphate, 4.7 μg/mL linoleic acid, 1× insulin–transferrin–selenium, 10 nM dexamethasone (all from Millipore Sigma, Burlington, MA, USA) and 10 ng/mL human TGF-β3 (PeproTech, Cranbury, NJ, USA).

### Biochemical assay to quantify ECM deposition

Samples were collected in 300 µL of passive lysis buffer (Promega, San Luis Obispo, CA, USA) and sonicated for 10 s at least 3 times. DNA content was quantified using the PicoGreen Quant-iT Assay Kit (Invitrogen, Waltham, MA, USA) according to manufacturer’s instructions. Sulfated GAG content was assessed using the DMMB assay. Total protein content was quantified using a BCA Protein Assay Kit (Thermo Fisher, Waltham, MA, USA) per the manufacturer’s instructions. Total collagen content was quantified using the Hydroxyproline Kit (Chondrex, Woodinville, WA, USA) per manufacturer’s instructions.

### Histological assessment

Samples were fixed in 4% paraformaldehyde (PFA) for 1 h, washed twice with PBS, embedded into Tissue-Tek OCT as described[31]. Samples were snap-frozen, sectioned at 8 μm on a cryostat, and mounted onto a charged slide. Samples were stained with Safranin-O, counterstained with Fast Green for GAG deposition using standard protocols, and imaged using an EVOS XL Core microscope. For immunohistochemistry, samples were rehydrated in PBS and then incubated in blocking buffer composed of 1x PBS, 0.5% Tween20, and 3% BSA for 1 h at room temperature. For detection of collagen type II, slides were incubated in collagen II polyclonal antibody (1:200, Invitrogen, AB_2818092) overnight at 4°C. Slides were then treated with a secondary goat-anti rabbit antibody conjugated with Alexa Fluor 647 (1:1000, Invitrogen, AB_2535812) for 1 h at room temperature. Slides were counterstained with DAPI (Thermo Fisher). Large tile scans were taken at 20x using confocal microscopy (Leica Stellaris 5, Wetzlar, Germany).

### Assessment of macrophage polarization

IC-21 murine macrophages (ATCC) were expanded in RPMI 1640 (ATCC) supplemented with 10% FBS. Macrophages were polarized toward an M1-like phenotype by exposure to lipopolysaccharide (LPS, 200 ng/mL, Thermo Fisher), while macrophages were polarized toward an M2-like phenotype by exposure to interleukin-4 (IL-4, 20 ng/mL, Thermo Fisher). LPS and IL-4 were added 48 h before seeding and refreshed with basal media after 24 h. The microgel slurry was mixed with macrophages at 8×10^6^ cells/mL and pipetted into a 5 mm x 1.5 mm cylindrical mold using a positive pressure displacement pipette and exposed to UV light for 90 s to form annealed microgel scaffolds. After 24 h, gels were moved to a new plate with fresh media and maintained in culture for 3 d with media changes every day. Cells were recovered from annealed scaffolds using ice cold 2.5 mM EDTA in PBS for 3 min and gently mixed, followed by dilution with basal media. Solutions were filtered through a 30 µm sieve to create a single cell suspension, and macrophage polarization was characterized using flow cytometry (Attune NxT, Life Tech, Waltham, MA, USA) as previously described[14]. Following Fcγ receptor blocking (1:40, TruStain FcX, BioLegend, San Diego, CA, USA), cells were stained with antibodies against F4/80 (1:50, eBioscience #MF48021, San Diego, CA, USA), CD86 (1:160, eBioscience #47-0862-82), and CD206 (1:40, eBioscience #48-2061-82). Cellular viability was evaluated with fixable Zombie Aqua (1:250, Life Tech, Carlsbad, CA, USA). Cells were then fixed with 2% PFA, permeabilized with 0.1% Triton-X, and stained for intracellular markers, iNOS (1:500, eBioscience #12-5920-82) and Arginase-1 (1:500, eBioscience #53-3697-82), overnight at 4°C with gentle agitation. An M1-like phenotype was characterized by F4/80+CD86+iNOS+ populations and M2-like phenotypes by F4/80+CD206+ARG1+ populations.

### Murine microfracture model

We examined the effect of acellular, annealed microgel scaffolds on articular cartilage regeneration following microfracture in eight-week-old skeletally mature C57BL/6 male mice (Jackson Laboratory, Bar Harbor, ME, USA). All experimental animal procedures were conducted in accordance with UC Davis Institutional Animal Care and Use Committee guidelines (Protocol #24191) and National Institute of Health animal handling standards. We performed a cartilage microfracture model as previously described[32]. Briefly, animals were anesthetized with isoflurane and maintained under general anesthesia delivered through a nose cone. A 5 mm incision was made medial to the knee, the patella was reflected laterally, and the knee was flexed to expose the femoral condyles. The defect was created using a 1 mm diameter awl to access the underlying cartilage surface. A 1 mm biopsy punch was used to collect and apply the scaffold directly to the microfracture defect site and retained in position by the surrounding anatomy. The sham control did not receive any implant or treatment. The patella was repositioned, and the incision was closed using a 6-0 Prolene suture. Animals were euthanized 8 weeks postoperatively, and femurs were harvested for mechanical testing and histology.

### Mechanical testing of murine articular cartilage

Murine femurs were harvested, cleaned, and stored in PBS at 4°C prior to testing. We measured the mechanical properties of articular cartilage using a Mach-1 micromechanical testing system (Biomomentum, Laval, QC, Canada). Before mechanical testing, samples were equilibrated in PBS at room temperature to prevent tissue dehydration throughout testing. Femurs were secured to a custom 3D-printed testing stage to maintain consistent positioning during indentation measurements. Cartilage mechanical properties were assessed using a normal indentation protocol with a spherical indenter tip (0.3 mm diameter) and a 1.5 N load cell. The indenter was lowered onto the cartilage surface at a Z-contact velocity of 0.05 mm/s. Indentation testing was performed to a depth corresponding to approximately 20% of the cartilage thickness using an indentation amplitude of 0.03 mm and an indentation velocity of 0.03 mm/s. Indentation measurements were collected from regions within the defect, along the defect edge, and in the surrounding native cartilage tissue, with multiple measurements obtained from each region per sample. The instantaneous modulus of the cartilage was calculated from the force-displacement response obtained during indentation testing. The normalized modulus was calculated using the equation below.

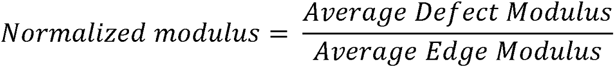

### Statistical Analysis

Statistical analysis was performed using GraphPad Prism 11. All data were tested for normal distribution *via* the Shapiro-Wilk normality test. Statistical significance was assessed by Student’s t-test, one-way ANOVA with Tukey’s multiple comparisons test, or two-way ANOVA. A p-value of less than 0.05 were considered statistically significant. Different letters denote statistical significance between groups, while data sharing a letter are not statistically different from one another.

## RESULTS

### Incorporation of azide and proteins do not alter microgel surface charge and compressive modulus

Microgels were designed to promote chondrogenesis through biophysical cues such as stiffness, surface charge, and local presentation of a chondroinductive growth factor. We incorporated an azide into the polymer to enable tethering of FGF18 *via* bioorthogonal click chemistry. We used DBCO-488 to confirm the presence of azide on the surface of the microgels and verify the reaction (**Figure 1A**). The conductivity of bulk hydrogels was measured using a previously published custom two-point probe setup. As expected, the nonionic polymer was less conductive than anionic and zwitterionic groups (**Figure 1B**). We measured the zeta potential of each monomer solution as an indicator of surface charge. We used bovine serum albumin (BSA) as a model protein and tethered it to the surface of microgel scaffolds. The incorporation of the azide and tethered-BSA did not alter the surface charge of microgel scaffolds (**Figure 1C**), confirming that persistence of the corresponding surface charge remained unchanged. Then, we quantified the electrostatic interaction of a cationic protein, lysozyme, onto annealed microgel scaffolds to test whether incorporation of azide and tethered-BSA affected protein adsorption onto the microgel surface. Lysozyme preferentially adsorbed to the anionic microgel scaffolds with minimal binding to the nonionic and zwitterionic groups regardless of azide or protein tethering (**Figure 1D**). We measured the release of tethered FGF18 in acellular microgel scaffolds at day 7. The release of FGF18 was negligible and did not differ across groups, confirming a stable bond (**Figure S1**). Collectively, these data demonstrate that azide modification of the polymer does not alter the biophysical properties of the microgel.

**Figure 1.**
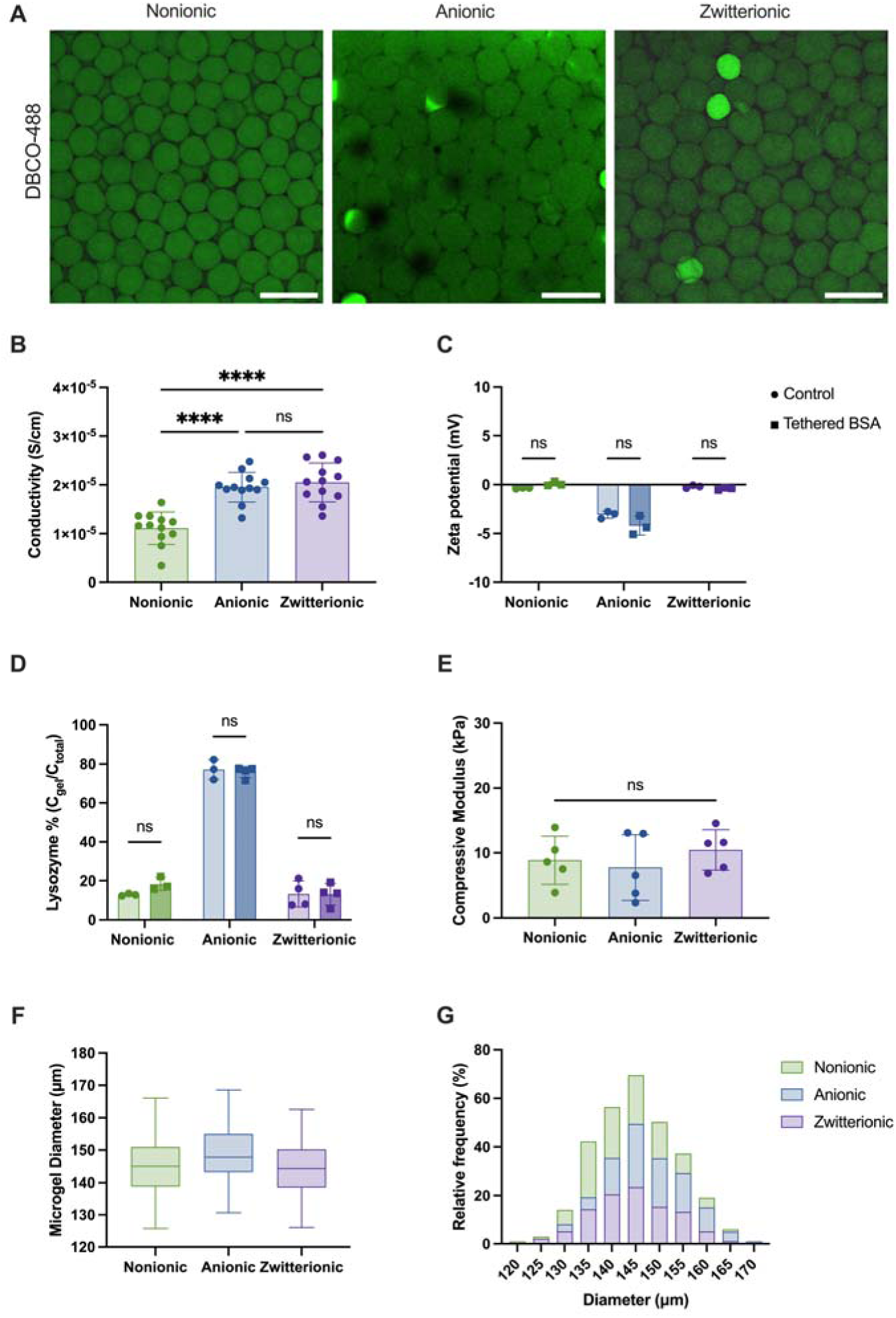
Surface charge and mechanical properties are preserved following azide functionalization and protein tethering. **(A)** Visual detection of DBCO-488 on surface of microgels confirms click reaction. Scale bar is 200 μm. **(B)** Nonionic polymer was less conductive than other polymer groups (n=12). **(C)** The incorporation of azide and tethered BSA did not alter zeta potential of polymer solutions (n=3). **(D)** Cationic lysozyme adsorbed more to anionic granular hydrogels at 24 h (n=3). **(E)** Microgel compressive modulus (n=5). **(F, G)** Microgel diameter across all groups (n=100). Significance determined by unpaired student t-test or one-way with multiple comparisons test (p<0.05). ns = non-significant.

Microgel compressive modulus remained consistent across groups, with an average compressive modulus of 9.0 kPa (**Figure 1E**). Specifically, nonionic, anionic, and zwitterionic microgels exhibited a compressive modulus of 8.9 ± 3.7 kPa, 7.8 ± 5.1 kPa, and 10.5 ± 3.1 kPa, respectively. Microgel diameter was quantified using ImageJ analysis and exhibited consistent average diameter of approximately 145 μm across all microgel groups (**Figure 1F, G**). Nonionic microgels possessed an average diameter of 145.3 ± 8.1 μm, anionic microgels were 147.8 ± 8.4 μm, and zwitterionic microgels had average diameter of 144.5 ± 8.4 μm.

### Tethered FGF18 promotes matrix accumulation in annealed microgel scaffolds

To evaluate the chondrogenic potential of microgels with varying surface charge and tethered FGF18, we seeded microgel scaffolds with mMSCs, annealed the construct with light, and cultured scaffolds in chondrogenic media. We compared these constructs to microgel scaffolds functionalized with an azide group but lacking FGF18 (control). After 3 weeks in culture, we observed visual differences in groups tethered with FGF18 (**Figure 2A**). Compared to control scaffolds of any charge, groups with tethered FGF18 had increased ECM matrix deposition evidenced by their darker color. Cells in nonionic and zwitterionic scaffolds exhibited more rounded and aggregated morphology while cells anionic scaffolds appeared more monodisperse (**Figure 2B**). We detected increases in GAG content in nonionic and anionic scaffolds tethered with FGF18 (**Figure 2C**) along with a significant increase in collagen in microgel scaffolds tethered with FGF18 (**Figure 2D**). Tissue maturation was assessed by measuring the compressive modulus of scaffolds following 3 weeks in culture (**Figure 2E**). Microgel scaffolds tethered with FGF18 exhibited higher moduli across all surface charges. We observed greater differences in compressive modulus for zwitterionic and anionic groups compared to nonionic groups. These data confirm that local and sustained presentation of FGF18 supports mMSC chondrogenesis and outperforms unmodified annealed microgel scaffolds.

**Figure 2.**
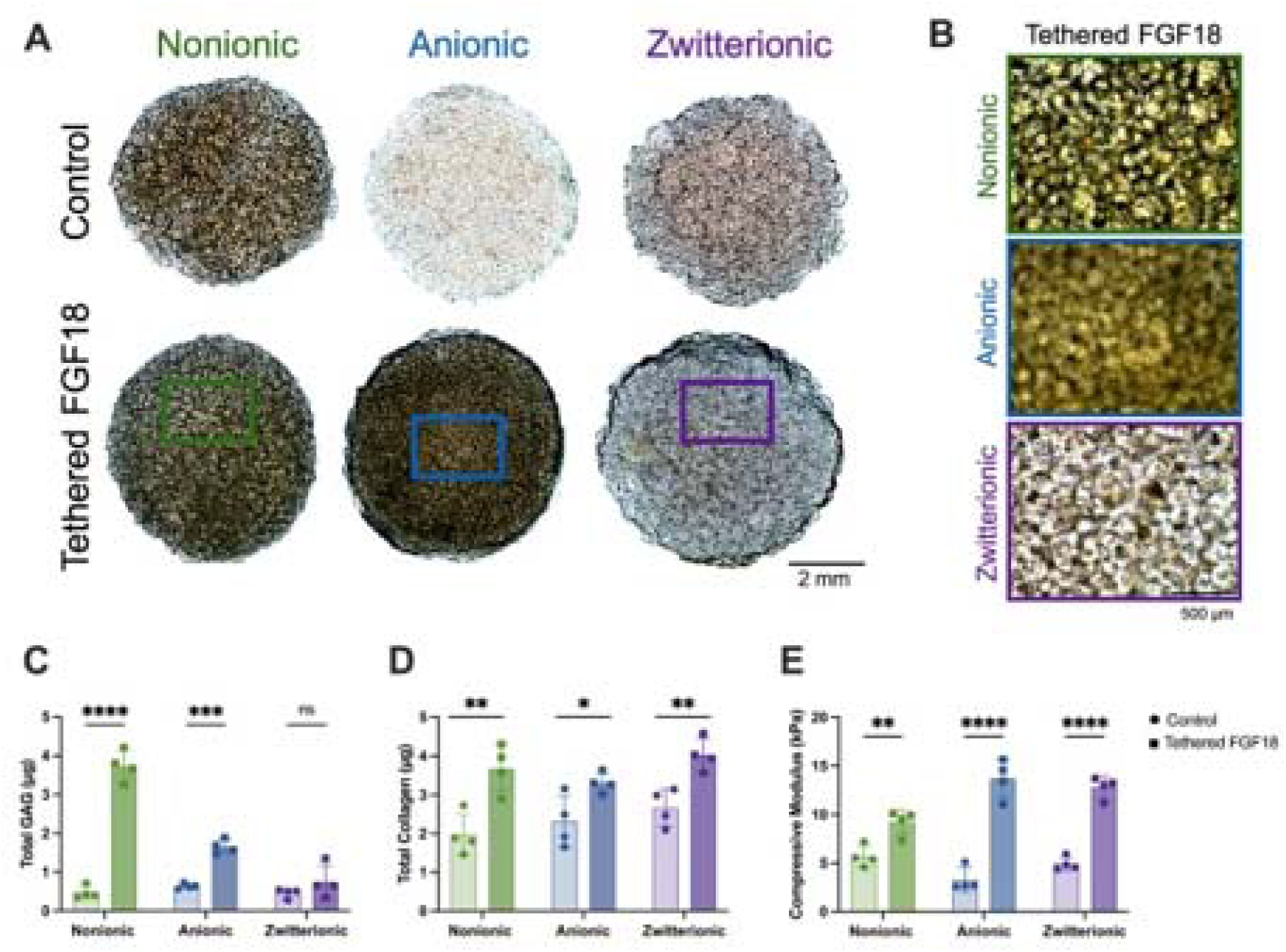
Tethered FGF18 promotes chondrogenesis in annealed microgel scaffolds. **(A, B)** Representative brightfield images of microgel scaffolds following 3 weeks in chondrogenic media. Scale bars are 2 mm and 500 μm. **(C)** GAG content, **(D)** collagen content, and **(E)** compressive moduli were higher in microgel scaffolds tethered with FGF18 (n=4). Significance determined by unpaired student t-test. ns = non-significant.

### Influence of surface charge and growth factor presentation on macrophage polarization

Macrophages play a key role in the inflammatory microenvironment of damaged cartilage. To investigate how surface charge and tethered FGF18 influence macrophage polarization, we cultured IC-21 murine macrophages in annealed microgel scaffolds with distinct surface charge. We then collected after 3 days to assess macrophage polarization *via* flow cytometry (**Figure 3A**). Cell viability was high within nonionic and zwitterionic groups (**Figure 3B**). In contrast, anionic scaffolds with tethered FGF18 exhibited reduced macrophage viability. We evaluated macrophage polarization by quantifying M1-like (CD86+iNOS+) and M2-like (CD86-CD206+ARG-1+) macrophages within the F4/80+ population. The number of M1-like macrophages were relatively similar across groups with an increased trend in FGF18-tethered scaffolds (**Figure 3C**). No significant differences were observed in the frequency of M2-like macrophages across groups (**Figure 3D**). Collectively, these findings demonstrate that surface charge and tethering of FGF18 has limited effects on macrophage polarization.

**Figure 3.**
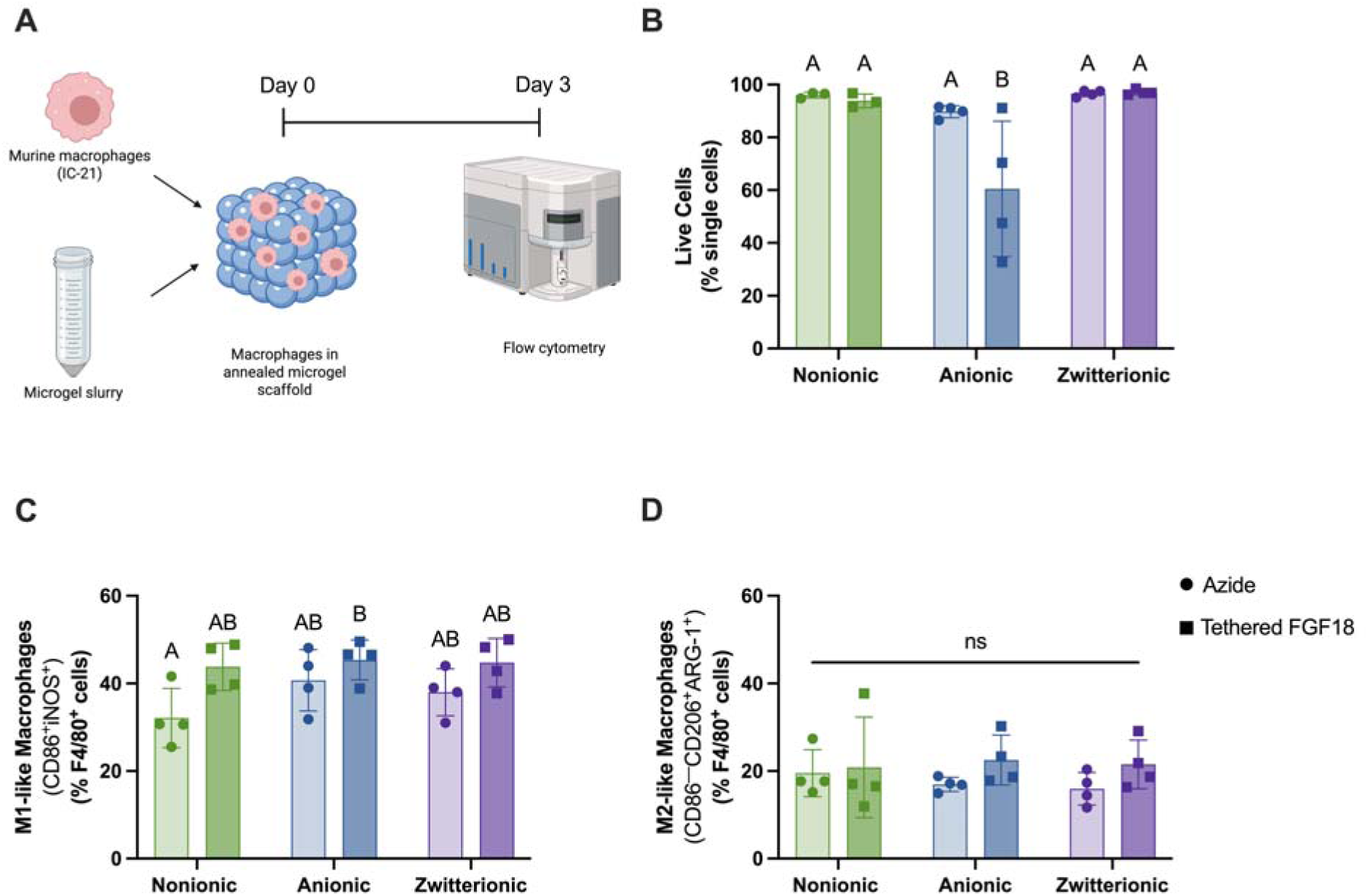
Surface charge and tethering of FGF18 has limited effects on macrophage polarization. **(A)** Schematic of *in vitro* studies to assess macrophage response to microgel scaffolds. **(B)** Percentage of live single cells across groups (n 3). **(C, D)** Quantification of M1-like and M2-like macrophages (n 3). Significance determined by two-way ANOVA with multiple comparisons test. Groups with different letters are significantly different (p<0.05). ns = non-significant.

### FGF18-tethering to microgel scaffolds promotes cartilage regeneration in murine MF model

We implanted acellular microgels scaffolds with distinct surface charges and functionalized with FGF18 to evaluate their regenerative potential in a murine microfracture (MF) model. Samples were collected after 8 weeks, and the mechanical properties of repair tissue were immediately assessed (**Figure 4A**). Measurements were performed within the defect, at the defect edge, and in the surrounding tissue as a reference point (**Figure 4B**). Representative heat maps revealed differences in instantaneous modulus across surface charges and due to FGF18 presentation (**Figure 4C**). We did not detect significant differences in average defect modulus across groups (**Figure 4D**). To account for variability across samples, we normalized the defect modulus to the defect edge modulus. In this case, full regeneration across the defect would be evidenced by a normalized modulus near 1, represented by the dashed line. Nonionic and zwitterionic scaffolds with FGF18 exhibited increased normalized modulus compared to their control, while normalized modulus in anionic groups was similar (**Figure 4E**). This is likely a result of the anionic control having higher normalized modulus compared to nonionic and zwitterionic groups.

**Figure 4.**
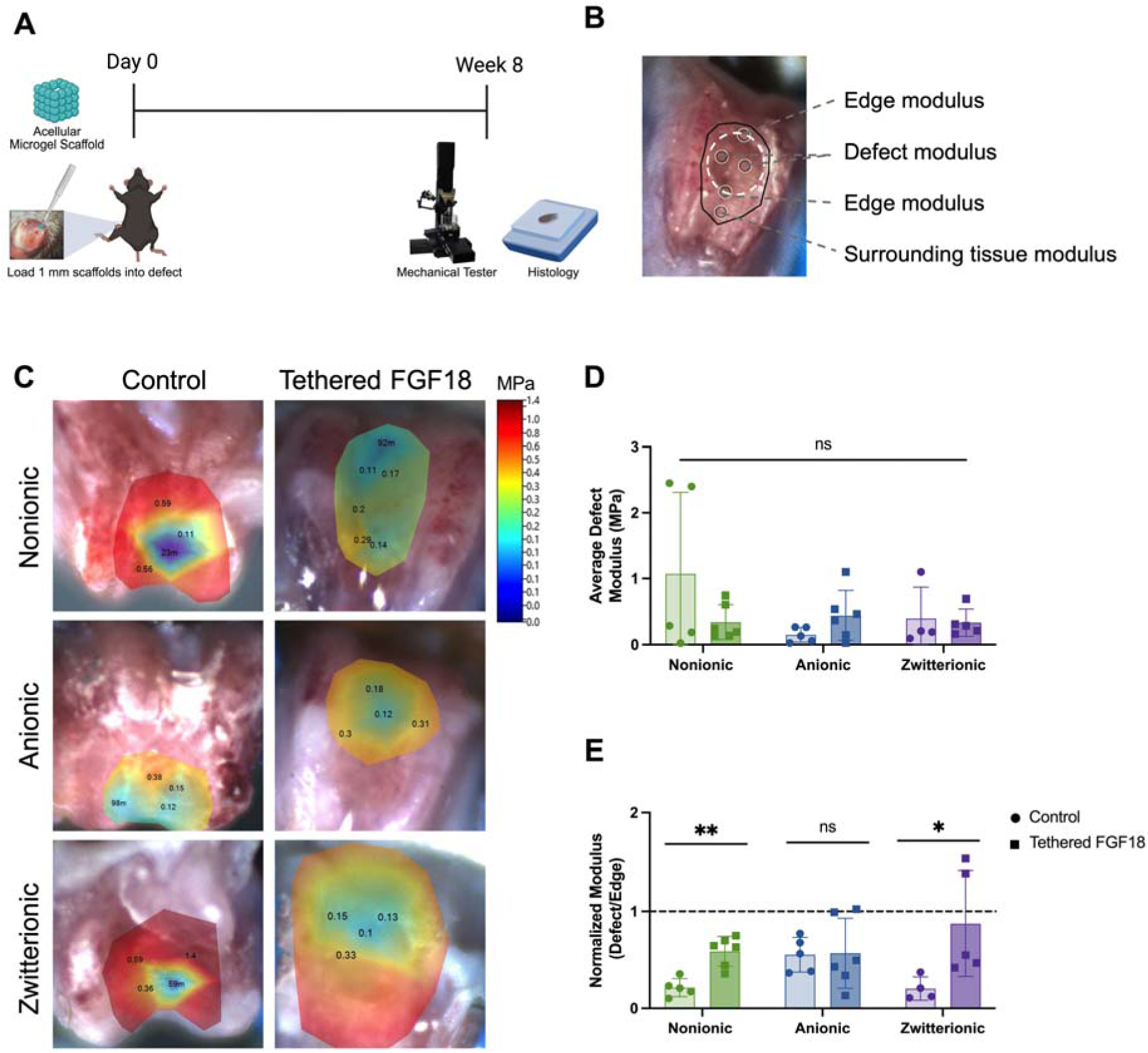
Tethered FGF18 increases normalized modulus of cartilage repair tissue in nonionic and zwitterionic scaffolds. **(A)** Acellular microgel scaffolds were implanted into a murine microfracture defect and evaluated after 8 weeks using mechanical testing. **(B)** Representative image of the cartilage defect region and mapping area used for mechanical analysis. **(C)** Representative heat maps of cartilage mechanical properties within the defect region and surrounding tissue. **(D)** Average instantaneous modulus of defect did not differ across groups (n 4). **(E)** Tethering of FGF18 increased normalized modulus in nonionic and zwitterionic groups (n 4). Dashed line defined as ratio where defect modulus = edge modulus. Significance determined by Student’s t-test with multiple comparisons post-hoc test (p<0.05). ns = non-significant.

Following mechanical testing, samples were processed and stained for histological analysis. Safranin-O staining confirmed sulfated GAG deposition within the defect (**Figure 5A**). We observed qualitative differences in GAG deposition in anionic and zwitterionic microgel scaffolds tethered with FGF18. The composition of MF defects was quantified using ImageJ (**Figure 5B**). Anionic and zwitterionic microgel scaffolds tethered with FGF18 exhibited 40% cartilage regeneration, on average. However, only defects treated with zwitterionic microgels tethered with FGF18 resulted in significantly higher cartilage regeneration compared to its control (**Figure 5C**). We subsequently examined the distribution of collagen type II as a marker of chondrogenesis (**Figure 6A**), and we observed an increase of collagen type II in groups tethered with FGF18. Collagen II was more prevalent in anionic and zwitterionic microgel scaffolds tethered with FGF18 (**Figure 6B**). Collectively, these data highlight that local sustained delivery of FGF18 supports *in vitro* and *in vivo* chondrogenesis and can be used synergistically with surface charge.

**Figure 5.**
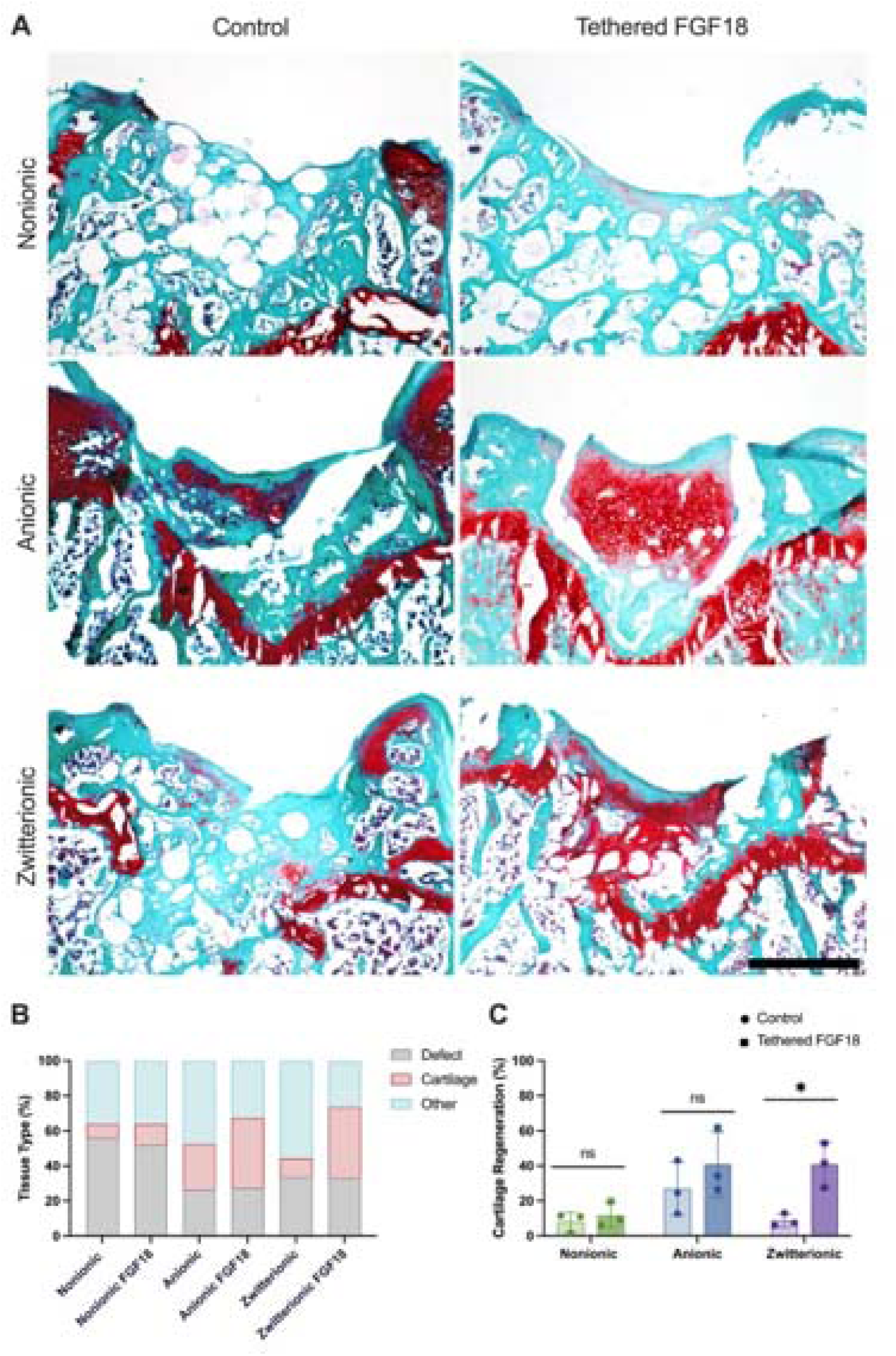
Tethered FGF18 enhances cartilage regeneration in MF model. **(A)** Representative images of Safranin-O staining after 8 weeks in MF defect. Scale bar is 500 μm. **(B)** Quantification of composition of defect (n=3). **(C)** Percentage of cartilage regeneration within defect. Significance determined by student t-test with multiple comparisons test (n=3 per group, *p<0.05). ns = non-significant.

**Figure 6.**
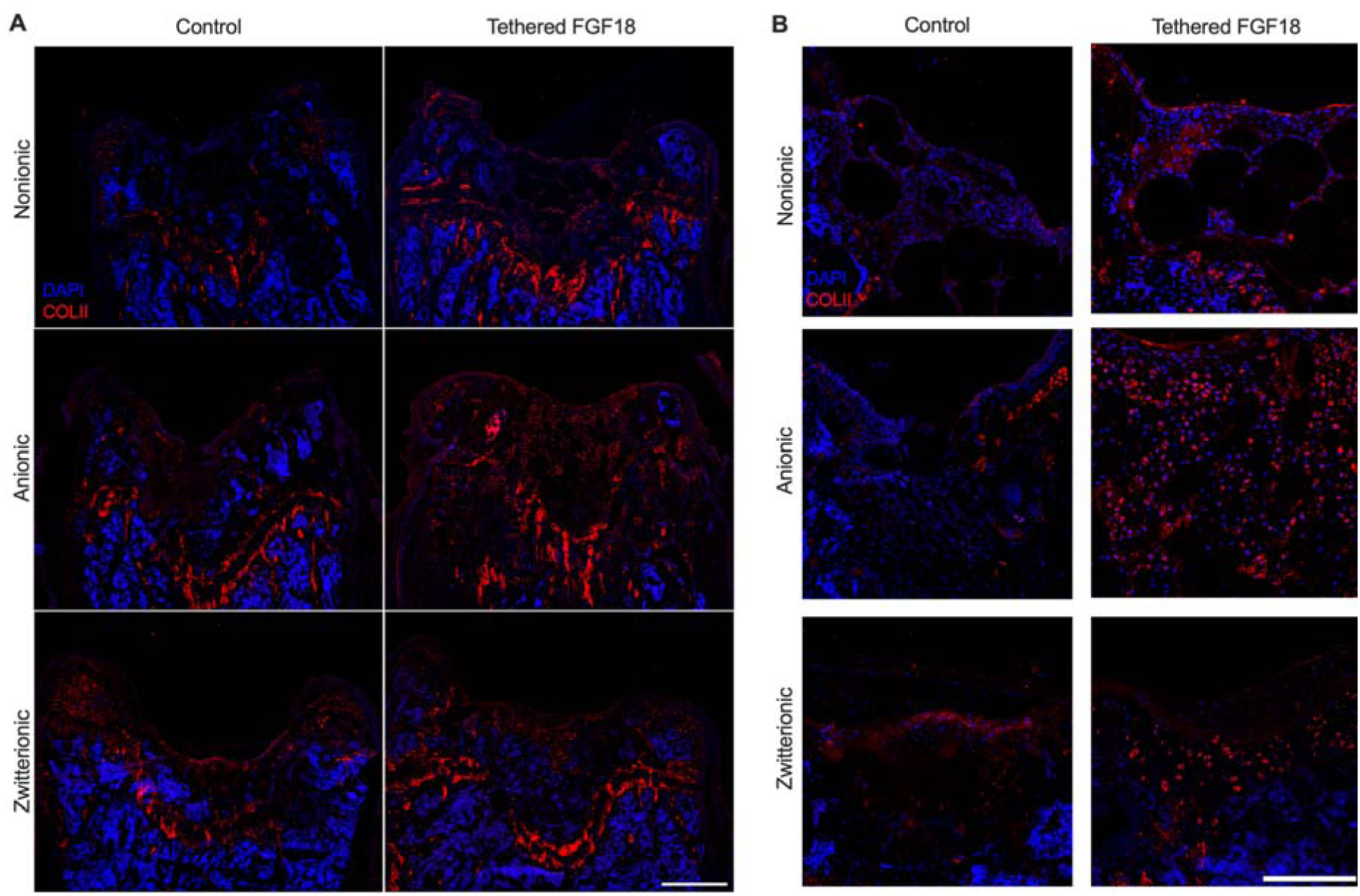
Tethered FGF18 promotes Collagen type II protein expression. (A) Representative tilescans at 20x magnification of collagen type II. Scale bar is 500 μm. (B) Representative images at 20x magnification demonstrate collagen II staining is increased in groups tethered with FGF18. Scale bar is 200 μm.

## DISCUSSION

Articular cartilage has limited self-healing capacity, requiring therapeutic options to stimulate repair. As one approach, engineered scaffolds are under development to recapitulate the mechanical and biological complexity of native cartilage tissue. Natural and synthetic polymers, including hyaluronic acid, collagen, and PEG, have been widely explored for cartilage regeneration, with crosslink density, water content, degradation, and surface chemistry being critical determinants of chondrogenic ECM production[15, 33, 34]. Annealed microgel scaffolds have emerged as a promising platform to overcome this challenge, as their inherent macroporosity facilitates rapid cell infiltration, nutrient transport, and ECM remodeling[35]. Their modularity allows for independent tuning of stiffness, porosity, and surface chemistry, making them well-suited for supporting chondrogenesis. Building upon prior work demonstrating that zwitterionic microgel scaffolds support ECM accumulation[24, 25], this study tests whether presentation of FGF18, a recombinant growth factor with prior clinical evidence of preserving cartilage thickness, from microgels with distinct surface charge can further enhance chondrogenesis. FGF18 was selected for its role in promoting chondrogenesis while suppressing hypertrophic cartilage progression[36, 37]. To our knowledge, this is the first study to investigate the interplay of localized growth factor delivery within microgel scaffolds with distinct surface charges for cartilage tissue regeneration. Furthermore, we demonstrate that azide functionalization enables stable growth factor delivery without compromising scaffold properties, and FGF18 promotes chondrogenesis and cartilage regeneration *in vivo*.

We covalently conjugated FGF18 to the azide-functionalized microgel surface *via* bioorthogonal click chemistry, providing a stable and spatially controlled strategy for growth factor presentation. Given that surface modifications can alter material properties critical for chondrogenesis, we measured zeta potential and conductivity to confirm that functionalization did not alter these properties. Bovine serum albumin (BSA) was used as a model protein that did not significantly alter surface charge or protein adsorption compared to their respective controls. Monodisperse microgels with an average diameter of ∼145 μm possessed comparable compressive moduli of ∼9 kPa across microgel formulations, establishing the capacity of this approach to decouple the effect of material stiffness, porosity, and surface charge. Previous studies have established that 9 kPa is an intermediate stiffness known to promote chondrogenesis[38, 39]. By matching microgel diameter and compressive modulus, any observed differences in chondrogenic outcomes may be attributed to surface charge or FGF18 signaling.

We investigated the influence of surface charge and localized FGF18 delivery on chondrogenesis by seeding annealed microgel scaffolds with mMSCs. All microgel groups were functionalized with RGD peptide to promote cell adhesion and an azide to facilitate protein tethering. Qualitative assessment revealed greater ECM deposition in tethered FGF18 scaffolds relative to controls. Anionic scaffolds exhibited higher cell density with more monodisperse cell distribution, whereas nonionic and zwitterionic scaffolds displayed rounded, aggregated cell morphology consistent with chondrogenic differentiation. When compared to microgel scaffolds of the same stiffness and surface charge, microgels tethered with FGF18 possessed greater GAG content and increased compressive modulus. These results are consistent with prior work demonstrating that covalent immobilization of growth factors within scaffolds outperforms soluble delivery or physical entrapment in promoting ECM production[40, 41]. Collectively, these data demonstrate that tethering of FGF18 to microgel scaffolds enhances chondrogenic differentiation, outperforming unmodified microgel scaffolds. Previous studies have tethered TGF-β[40, 42] and N-cadherin mimetic peptides [11, 30, 43] to scaffolds to drive chondrogenesis. While these approaches effectively induce chondrogenic commitment, FGF18 promotes chondrogenic differentiation while also enhancing ECM production and cartilage-specific tissue formation, resulting in a mechanistically distinct approach for cartilage regeneration.

To evaluate the translational potential of this microgel platform, we investigated cartilage regeneration using an established murine microfracture model[32]. This model provided clinically relevant context for examining the combined effects of surface charge and local FGF18 delivery on functional tissue repair. Mechanical testing revealed that defects treated with nonionic and zwitterionic scaffolds with tethered FGF18 exhibited significantly higher instantaneous moduli compared to their respective controls. The increase in normalized moduli is attributed to enhanced ECM deposition and tissue integration within the scaffold and surrounding defect. These findings are consistent with our *in vitro* chondrogenesis data, collectively supporting the hypothesis that sustained localized FGF18 delivery promotes restoration of mechanical properties relevant to load-bearing function.

Histological assessment of tissue formation, GAG deposition, and collagen type II staining highlighted the potency of FGF18 in driving cartilage regeneration. In the absence of FGF18, microgel scaffolds had remarkably less cell infiltration and greater void space within the defect. When evaluating surface charge alone, anionic and zwitterionic scaffolds increased cartilage regeneration compared to nonionic scaffolds. However, tethered FGF18 substantially improved cartilage regeneration beyond what surface charge alone could achieve. Anionic and zwitterionic scaffolds tethered with FGF18 demonstrated the greatest cartilage regeneration with increased cell infiltration and matrix deposition. Although our *in vitro* studies suggested that nonionic microgels could support chondrogenic differentiation, these scaffolds resulted in limited tissue regeneration *in vivo*. This discrepancy likely reflects the potent instructive cues within the chondrogenic media that promote chondrogenesis independent of scaffold-mediated signaling, whereas tissue regeneration in the MF model relies heavily on endogenous cell infiltration, matrix remodeling, and endogenous biochemical cues. Overall, these findings support FGF18 as a potent stimulus for cartilage regeneration, capable of enhancing outcomes independent of scaffold surface charge.

While this study provides meaningful insight into how surface charge and localized growth factor delivery influence cartilage regeneration, several limitations must be acknowledged. The use of young, healthy mice does not fully recapitulate the degenerative microenvironment of OA, in which chronic inflammation and impaired cellular function hinder regeneration. Future studies should consider aged mouse models or surgically induced OA such as destabilization of the medial meniscus (DMM), which more closely recapitulates clinical conditions[44]. Also, since the FORWARD trial demonstrated a dose dependent response, future studies should investigate the dose dependency in a sustained localized delivery compared to a soluble growth factor delivery[28]. Future iterations could incorporate additional chondrogenic cues such as TGF-β3, which has a synergistic effect when delivered with FGF18 to promote robust articular cartilage formation[45]. Lastly, the incorporation of a degradable crosslinker such as GPQ-A, a matrix metalloproteinase (MMP)-degradable crosslinker, may better support tissue remodeling upon implantation[17, 46]. Addressing these limitations would improve the translation of this platform towards clinically relevant cartilage repair strategies.

We engineered microgels with distinct surface charges and covalently linked FGF18 to investigate the synergistic effect of substrate charge and sustained growth factor presentation on cartilage regeneration. By using bioorthogonal click chemistry, FGF18 was covalently tethered to the microgel surface to enable localized and sustained presentation within the microgel scaffold. Annealed microgel scaffolds tethered with FGF18 enhanced extracellular matrix deposition, improved compressive mechanical properties, and promoted cartilage repair compared to their surface charge counterparts. These findings suggest that while surface charge contributes to regulating cell-material interactions, sustained presentation of FGF18 serves as a dominant anabolic cue driving chondrogenic regeneration. Overall, this study highlights the therapeutic potential of combining annealed microgel scaffolds with tethered FGF18 to create instructive biomaterial platform for functional cartilage repair.

## Generative AI disclosure statement

During the preparation of this work, the authors used ChatGPT to improve readability and grammar. After using this tool/service, the authors reviewed and edited the content as needed and take full responsibility for the content of the published article.

## Data availability

The data that support the findings of this study are available from the corresponding author upon reasonable request.

## Acknowledgements

This work was supported in part by the National Institutes of Health (grant number R01 AR079211 to JKL). This manuscript is the result of funding in whole or in part by the National Institutes of Health (NIH). It is subject to the NIH Public Access Policy. Through acceptance of this federal funding, NIH has been given a right to make this manuscript publicly available in PubMed Central upon the Official Date of Publication, as defined by NIH. JKL gratefully acknowledges financial support from the Lawrence J. Ellison Endowed Chair of Musculoskeletal Research. Some figures were created with the use of Biorender.

## Author Contributions

**E.E.W**: Conceptualization, Methodology, Validation, Formal Analysis, Investigation, Writing – manuscript composition and editing. **H.-J.J.**: Investigation, Validation, Formal Analysis, Writing – editing. **K.C.W.**: Investigation, Validation, Formal Analysis. **K.C.**: Methodology, Investigation, Validation, Formal Analysis. **Y.W.**: Methodology, Investigation, Validation. **K.H.G.**: Methodology, Investigation, Validation, Formal Analysis **T.H.A.**: Conceptualization, supervision. **J.K.L.**: Conceptualization, supervision, writing - editing, financial support.

