## Supplementary material for "Bioorthogonal Tethering of FGF18 to Annealed Microgel Scaffolds Enhances Cartilage Repair": Figure S1

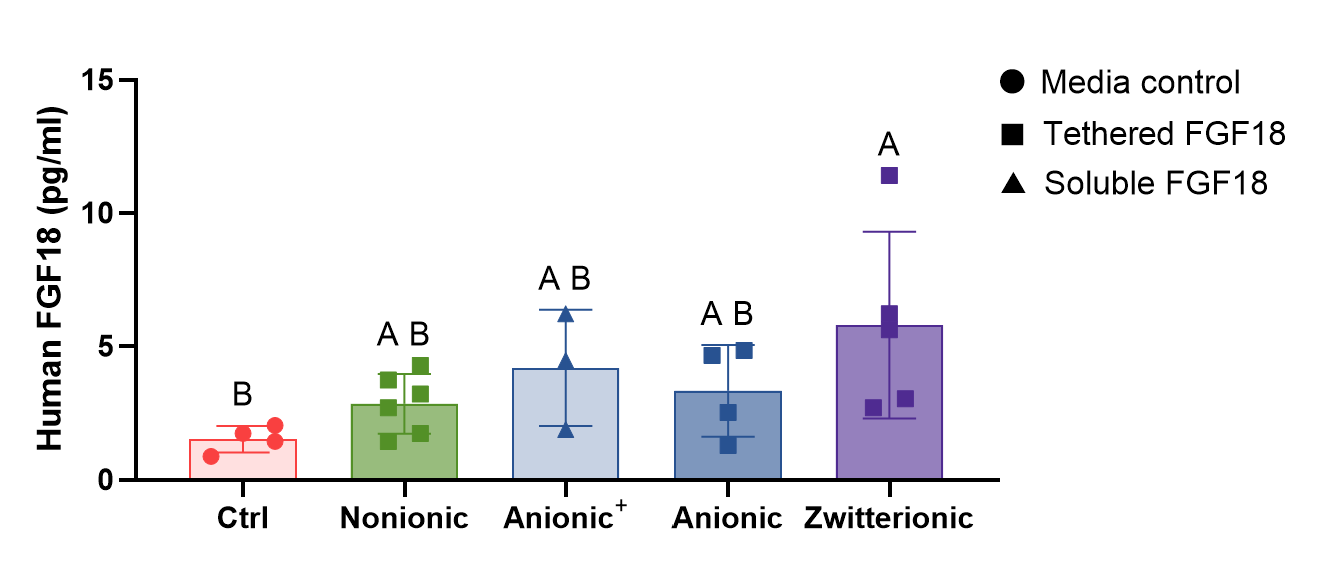

**Figure S1. Human FGF18 successfully immobilized to the microgel surface.** Release of FGF18 in acellular microgel scaffolds did not differ across groups at Day 7. Data represent mean ± SD (n=3-6). Significance determined by two-way ANOVA with multiple comparisons test. Groups with different letters are significantly different (p<0.05).
